# TFF3 is a ligand for LINGO2 that de-represses EGFR to control disease outcome during colitis and gastrointestinal nematode infection

**DOI:** 10.1101/469700

**Authors:** Yingbiao Ji, Yun Wei, JoonHyung Park, Li Yin Hung, Tanner Young, Karl Herbine, Taylor Oniskey, Christopher Pastore, Wildaliz Nieves, Ma Somsouk, De’Broski R. Herbert

## Abstract

Intestinal epithelial cells (IEC) comprise diverse lineages that serve distinct roles necessary for regulation of nutrient absorption, regeneration, immunity, and homeostasis^1,2^. Goblet cells secrete Trefoil factor 3 (TFF3) to maintain mucus viscosity and drive mucosal healing by inhibiting cell death and influencing tight junction protein expression^3^. However, whether TFF3 signaling relies upon conventional ligand-receptor interactions has been unclear for decades. This study demonstrates that the orphan transmembrane protein leucine rich repeat receptor and nogo-interacting protein 2 (LINGO2) immunoprecipitates with TFF3, that LINGO2 and TFF3 co-localize at the IEC cell surface, and that TFF3/LINGO2 interactions block IEC apoptosis. Loss of function studies show that TFF3-driven STAT3 and EGFR activation are both LINGO2 dependent. Importantly, we demonstrate that TFF3 disrupts LINGO2/EGFR interactions that normally restrict EGFR activity, resulting in enhanced EGFR signaling. Excessive EGFR activation in *Lingo*2 gene deficient mice exacerbates colitic disease and accelerates host resistance to parasitic nematodes, whereas TFF3 deficiency results in host susceptibility. Thus, our data demonstrating that TFF3 functions through a previously unrecognized ligand-receptor interaction with LINGO2 to de-repress LINGO2-dependent inhibition of EGFR activation provides a novel conceptual framework explaining how TFF3-mediates mucosal wound healing through enhanced activation of the EGFR pathway.

---

Although Trefoil factor 3 (TFF3) is well known to drive reparative pathways at respiratory, ocular, genitourinary and gastrointestinal mucosa, the identity of a potential TFF3 receptor has remained elusive for several decades ^4-8^. In addition to enhancing mucous viscosity ^9^, TFF3 induces epithelial cell survival and proliferation, activates EGFR, β-catenin, and STAT3 signaling pathways, and controls tight junction protein expression through ill-defined mechanisms to protect against gastrointestinal (GI) tissue injury and inflammation ^3,10^. Since the anti-inflammatory activity of TFF3 suggested that leukocytes may be directly responsive to TFF3, we utilized a human macrophage/monocyte cell line (U937) to first ask whether cytokine release could be regulated in response to rTFF3 exposure in order to isolate the TFF3 receptor. We found that rhTFF3 caused a dose-dependent reduction in endotoxin-mediated TNF release over a range of 1-100 ng/ml and that TFF3 suppression was lost at a higher range of rhTFF3 (100-1000 ng/ml) (Figure S1A). TFF3-induced interleukin 10 (IL-10) production was found over several orders of magnitude, but was lost at higher doses of rhTFF3 (Figure 1A). As the dose response is similar to the established range of activity for most cytokine and growth factor receptors ^11^, these observations suggested that TFF3 was interacting with a receptor in a conventional manner.

**Figure 1.**
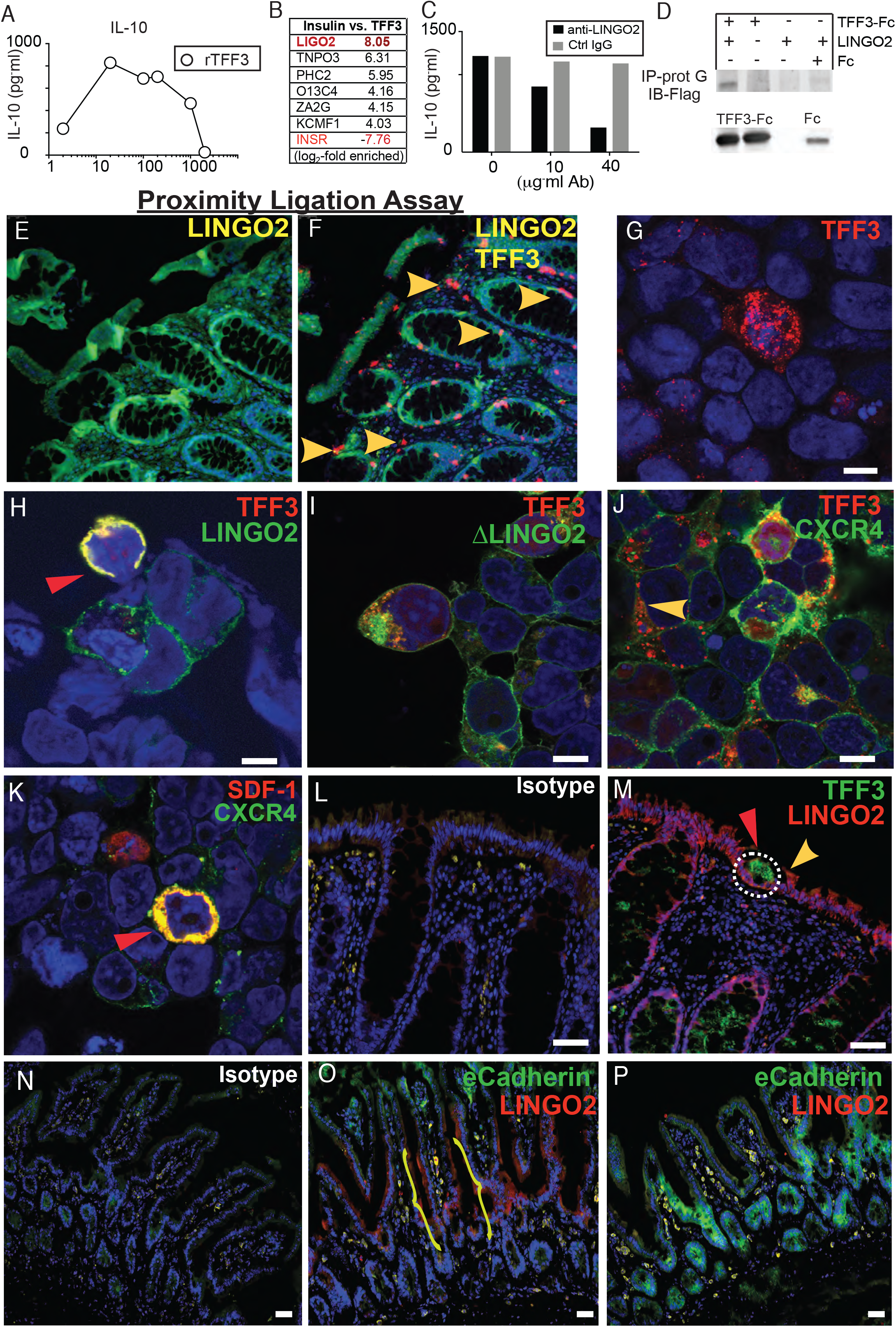
Identification of LINGO2 as a putative receptor for TFF3. (A) IL-10 production from the human macrophage cell line U937 in response to overnight exposure to different rhTFF3 concentrations. Representative of two independent experiments (B) Mass spectrometry fold-enrichment peptide score following TRICEPS screen of U937 cells exposed to rhTFF3 (C) IL-10 levels secreted by U937 production following overnight rTFF3 in the presence of anti-human LINGO2 mAb or isotype matched IgG. Representative of two independent experiments (D) Co-IP of TFF3-Fc and Lingo2:Flag expressed in in CHO cells IP was performed with protein G followed by immunoblotting with anti-Flag or anti-IgG. (E) Representative images from Proximity ligation assay performed on rectosigmoid biopsy samples from normal human subjects exposed to anti-LINGO2 or (F) anti-TFF3 and anti-LINGO2 mAb (G) Overexpression of HEK-293 cells single transfected with TFF3-RFP or (H) co-transfected with TFF3-RFP and LINGO2-GFP at 48 hr. (I) HEK-293 cells co-transfected with TFF3-RFP and NH_2_-terminal Flag-LINGO2 truncation mutant (Δ350 AA), (J) Overexpression of CXCR4-GFP and TFF3-RFP or (K) SDF-1-RFP on HEK-293 cells. (L) Representative immunofluorescence images of rectosigmoid biopsy samples from normal human subjects following co-staining with IgG isotype mAb or (M) anti-TFF3 mAb and anti-LINGO2 mAb. (O) Representative immunofluorescence images of ileum samples from wild-type or LINGO2KO C57BL/6 mice. Brackets indicate region of LINGO2 signal. Representative images are shown.

TFF3 was likely to interact with its receptor through low-affinity interactions because glycosylation has been shown critical for biological activity^12,13^, therefore U937 cells were subjected to the TRICEPS^TM^ protocol as a biochemical screening strategy to identify transmembrane protein(s) that bound TFF3 on the cell surface ^14,15^. As bait, rhTFF3 was covalently linked to the TRICEPS probe and incubated with PMA-treated U937 that had been gently oxidized with sodium periodate. Cell pellets were subjected to glycosidase digestion and peptides generated using mass spectrometry. To account for enrichment efficiency, recombinant human insulin was used as a bait to enrich for insulin receptor in the same experiment. Data show the TFF3-probe induced an 8-fold enrichment of LINGO2 (LIGO2) peptide, whereas the insulin-probe induced a 7.76-fold-enrichment in INSR peptide (Figure 1B), strongly suggesting that Leucine-rich repeat and immunoglobulin-like domain-containing nogo receptor-interacting protein 2 (LINGO2) was a potential receptor for TFF3.

LINGO2 is a 606aa protein with an extracellular domain (ectodomain), a transmembrane domain, and a cytoplasmic tail. The *LINGO2* gene is conserved between human and mouse, and belongs to the family of leucine-rich repeat and IgG-like domain proteins that includes LRIG1 and LRIG5 ^16^. In studies designed to further explore functional interactions between TFF3 and LINGO2, we found that anti-human LINGO2 Mab treatment of U937 resulted in a dose-dependent neutralization of TFF3-mediated TNF suppression (Figure S1b) and blocked TFF3-induced IL-10 production (Figure 1C). Additionally, co-immunoprecipitation experiments utilizing murine TFF3-Fc vs. empty vector Fc expressed and affinity purified from Chinese hamster ovary (CHO) cells revealed that TFF3 could pull down LINGO2 protein. Specifically, cell lysates from Flag-LINGO2 transfected CHO cells (600ug lysate) were incubated with either TFF3-Fc (20ug) or empty-Fc revealing that TFF3-Fc, but not Fc, co-precipitated Flag-LINGO2 (Figure 1D), further indicating that TFF3 directly bound LINGO2.

Given that epithelial cell lineages are considered the major source and target of TFF3 activity, two approaches were used to test whether LINGO2 captured TFF3 at the cell surface of IECs. A proximity ligation assay (PLA) on human rectal biopsy tissue was used to test whether the TFF3/LINGO2 interaction was within the range of a ligand receptor pair in a physiologically relevant context (<40nm), in a system that lacked genetic manipulation or overexpression ^17^. Compared to the lack of fluorescence signal with anti-LINGO2 mAb, combined staining with anti-TFF3 and anti-LINGO2 Ab revealed robust puncta within both the intestinal lamina propria and crypt epithelia (Figure 1E-F). As an alternative approach, HEK293 cells were double transiently transfected with constructs encoding GFP-LINGO2 and RFP-TFF3, fixed, permeabilized and imaged using confocal microscopy. Whereas RFP-TFF3 single transfectants had multiple fluorescent puncta throughout the cytoplasm of the positive cells (Figure 1G), 99% of the GFP-LINGO2/ RFP-TFF3 double-transfectants had clear yellow signal at the cell membrane, indicative of co-localization (Figure 1H and Figure S1C). To confirm the LINGO2 extracellular domain was necessary for the TFF3 interaction, we co-transfected TFF3-RFP with a NH_2_-terminal Flag-LINGO2 truncation mutant (Δ350 AA), which abrogated co-localization and resulted in TFF3 red fluorescent puncta, similar to single transfectants (Figure 1I). These data supported the accepted notion that the leucine-rich-repeat (LRR) region is responsible for ligand binding for the receptor kinases with leucine-rich-repeat ectodomains across the plant and animal kingdoms. To address reports suggesting that TFF3 may also signal through CXCR4, the canonical receptor for stromal derived factor-1 (SDF-1)^18^, CXCR4-GFP was co-transfected with TFF3-RFP or SDF1-RFP. We found that TFF3-RFP/CXCR4-GFP transfectants had red puncta, whereas 99% of SDF1-RFP/CXCR4-GFP double transfectants demonstrated clear evidence of co-localization via yellow signal (Figure 1J-K and Figure S1C). Taken together these data further indicated that TFF3 bound LINGO2.

Genome wide association study (GWAS) data show *Lingo2* genetic polymorphisms in human patients with cancer ^19^, asthma ^20^, chronic obstructive pulmonary disease ^21^, Parkinson’s ^22^, and Inflammatory bowel disease ^23^. Therefore, to understand the spatial pattern of TFF3 and LINGO2 interactions that might be relevant to some of these disease states, human rectal tissues were immunostained with fluorescent antibodies against TFF3 and LINGO2. Compared to isotype matched control staining, both enterocytes and crypt epithelia express LINGO2 whereas goblet cells located within nascent crypts were highly positive for TFF3 staining (Figure 1L-M and Figure S1D) and, in some instances, patches of cell-free TFF3 localized to the apical surface of LINGO2 positive enterocytes (Figure 1M). Next, we generated LINGO2 knockout mice (LINGO2KO) through non-homologous end joining (NHEJ) using CRISPR/CAS9 gene-editing to determine where LINGO2 was expressed in mice and whether gene deletion would have any biological impact on GI diseases. We created several LINGO2KO founder lines and selected one strain with a 200 bp deletion immediately downstream of the transcriptional start site (Figure S1E). As an initial characterization, intestinal tissues subjected to LINGO2 immunostaining revealed the expected absence of LINGO2 on the apical side of small intestinal villi in LINGO2KO mice compared to WT C57BL/6 controls (Figure 1 N-P). LINGO2 mRNA transcripts were detected in CD4, CD11c, and EpCAM positive cells from the intestines of WT mice, but largely absent in LINGO2KO mice (Figure S2A). Collectively, these data suggested a model where goblet cell-secreted TFF3 bound enterocyte-expressed LINGO2 to mediate biological functions through cognate ligand-receptor interactions on potentially different cell types.

TFF3-mediated cytoprotective functions are well-documented and are associated with activation of EGFR and ERK in various cell lines derived from pancreas, lung, and colon ^3,24,25^ ^26^. While TFF3 can regulate the transcription factor STAT3 by promoting Stat3 phosphorylation and *STAT3* transcriptional activation of TFF3 and other STAT3-dependent genes ^27,28^, these effects are not mediated by direct binding of TFF3 to the EGFR ^29^. To determine if TFF3 instead impacts EGFR signaling and subsequent STAT3-dependent signaling through LINGO2, we used a murine intestinal adenocarcinoma cell line (MC38) that expresses all four LINGO family member genes, with LINGO2 showing highest expression (Figure 2A). A LINGO2KO subclone of the MC38 cell line (Δ-LINGO2) was generated using CRISPR-mediated insertion of a GFP-H2-K^k^ knockin construct followed by FACS sorting (Figure 2B) showing a 20-fold reduction in LINGO2 expression (Figure 2C). To directly test whether TFF3 protected against cytotoxicity in a LINGO2-dependent manner, we compared the parental MC38 vs. Δ-LINGO2 MC38 lines in cell-death assays using the apoptotic agent, staurosporine. Whereas TFF3-Fc pre-treatment significantly reduced cytotoxicity of the parental MC38, this protection was abrogated in the Δ-LINGO2 line (Figure 2D). Given the anti-apoptotic function of STAT3, we then asked whether TFF3 driven STAT3 activation was LINGO2 dependent. Notably, TFF3-Fc (1ug/ml) induced STAT3 phosphorylation (pStat3 705) within 5 min in the parental MC38 line with a signal that slowly dissipated over 30 min, whereas TFF3-Fc-treated Δ-LINGO2 cells failed to up-regulate pSTAT3 to a similar extent (0.5 fold after 5 min treatment) and showed evidence of reduced total STAT3 levels (Figure 2E and 2G). Similarly, as TFF3 induces EGFR phosphorylation (pEGFR), we found that TFF3-Fc induced marked increase in pEGFR at residue 1068 within parental MC38, but TFF3-Fc only marginally increased pEGFR in Δ-LINGO2 cells (Figure 2F and 2G). Thus, LINGO2 was requisite for TFF3 to protect against apoptosis and for TFF3 driven STAT3 and EGFR activation.

**Figure 2.**
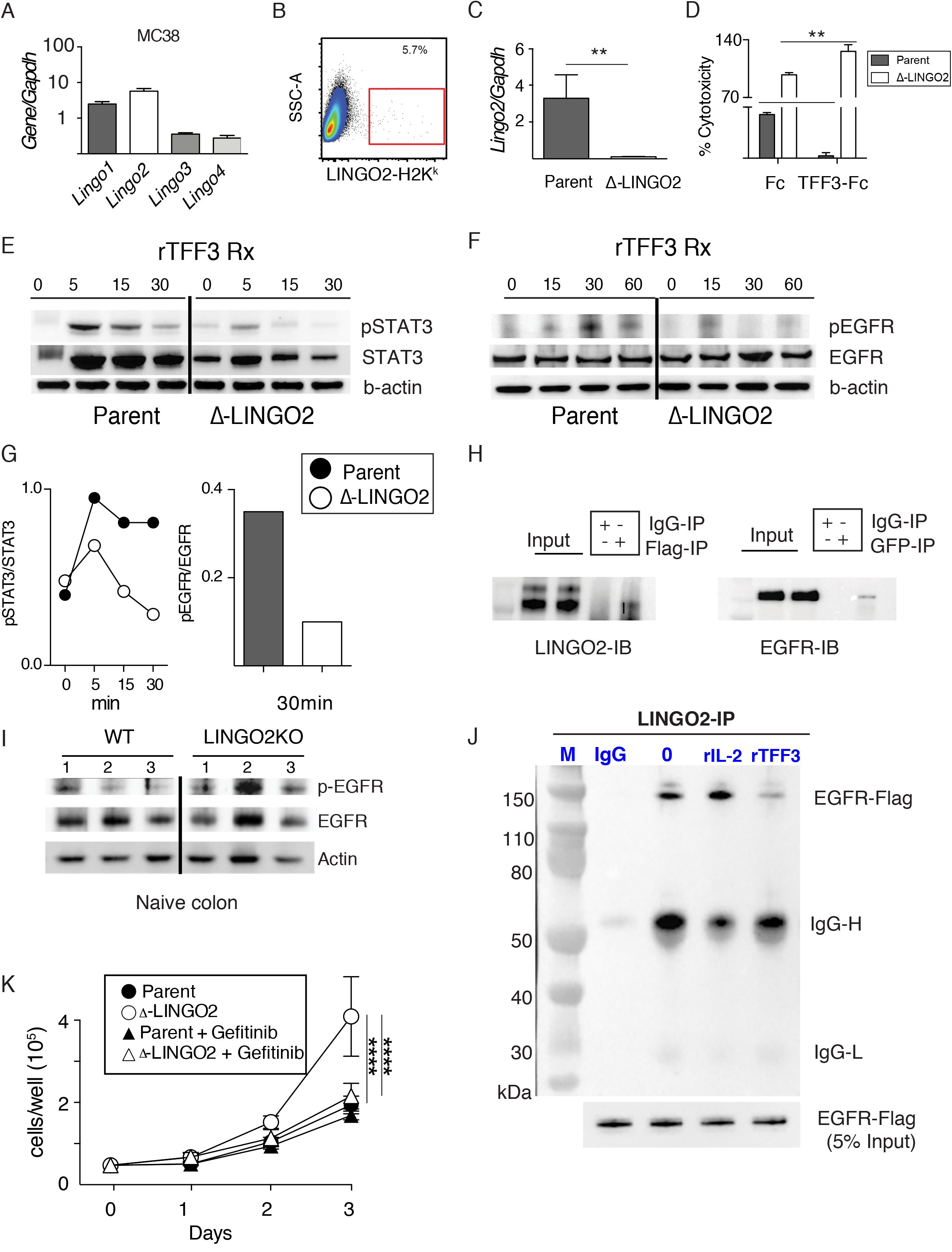
TFF3 requires LINGO2 to block cytotoxicity, regulate STAT3 and de-repress EGFR activation. (A) LINGO family mRNA transcript levels in the MC38 mouse intestinal cell line. Means ± SE of 3-6 replicates are shown. (B) Gating strategy for FACS-sorting of gene-edited MC38 cells via incorporation of the LINGO2-H2K^K^ knockin construct cell line designated Δ-LINGO2 (C) mRNA transcript levels for Lingo2 in the Δ-LINGO2 MC38 cell line quantified by RT-PCR (D) Staurosporine-induced cytotoxicity at 6 hr.-post exposure in MC38 vs. Δ-LINGO2 cells following overnight incubation in serum-free (1% FBS) media with Fc (1μg/mL), TFF3-Fc (1μg/mL or 23.81 nM) treatment. (E) Time-course of phospo-STAT3 and total STAT3 levels in parental MC38 vs. Δ-LINGO2 MC38 cells following exposure to TFF3-Fc (1μg/mL) and phospo-EGFR (Y-1068) vs. total EGFR levels compared to a beta-actin loading control (G) Densitometry measurements for data shown in “E” and “F” at 30 min (H) co-immunoprecipitation of LINGO2-Flag and EGFR-GFP in co-transfected HEK293 cells. IP was performed with anti-Flag followed by IB with anti-GFP (left) or IP with anti-GFP followed by IB with anti-Flag (right). (I) Western blot showing phospo-EGFR (Y-1068) vs. total EGFR levels compared to a beta-actin loading control in small intestinal lysates from individual WT vs. LINGO2KO mice. Each lane is an individual mouse (K) Co-IP of LINGO2-Flag and EGFR-GFP in HEK293 cell line in the presence of rTFF3 or rIL-2. (L) Proliferation curve over time for parental MC38 vs. Δ-LINGO2 MC38 with or without Gefitinib (5uM) treatment. Means ± SE of 6 replicates are shown.*=p<0.05 ****p<0.001

LINGO1 (a paralog of LINGO2) is a component of a heterotrimeric signaling complex comprised of the Nogo-66 receptor and the p75 neurotrophin receptor that inhibits axon regeneration ^30^. Interestingly, LINGO1 overexpression inhibits EGF-induced EGFR signaling ^31^, suggesting that LINGO1 blocks ligand-dependent EGFR activation. Notably, data from reciprocal co-immunoprecipitation experiments demonstrated that Flag-LINGO2 and GFP-EGFR interact (Figure 2H) and EGFR-GFP and LINGO2-RFP co-localize at the cell surface of co-transfected HEK293 cells (Figure S2B). Because TFF3 required LINGO2 to drive phosphorylation of EGFR (Figure 2F), we surmised that LINGO2 and EGFR were constitutively bound at the steady-state and that TFF3 binding to LINGO2 may sequester it from EGFR, thereby de-repressing EGFR activation by ligand-dependent or independent mechanisms. Consistent with this hypothesis, expression levels of EGFR-responsive genes EGF, HB-EGF, and Areg were increased in Δ-LINGO2 MC38 relative to the parental MC38 (Figure S2C). This was accompanied by a 3-fold higher sensitivity of Δ-LINGO2 MC38 cells to EGF-induced EGFR activation in comparison to the parental MC38 (Figure S2D), supporting the hypothesis that LINGO2 normally inhibits EGFR and suggesting that binding of TFF3 to LINGO2 could disrupt this inhibition, allowing EGFR to be more readily activated by its cognate ligands.

To test the ability of TFF3 to de-repress LINGO2-dependent EGFR inhibition we first obtained whole colonic tissues from LINGO2KO and WT mice and comparatively evaluated total vs. phosphorylated EGFR levels under steady-state conditions. Consistent with our prediction, LINGO2KO colonic tissue had higher basal levels of pEGFR compared to WT tissue (Figure 2I). Consistent with enhanced EGFR activity, quantification of S-phase crypt epithelial cells under steady state conditions using co-immunostaining with anti-E-cadherin and anti-Ki-67 mAb revealed a higher basal number of Ki-67 positive cells in the LINGO2KO compared to the WT (Figure S2E) Second, we devised a strategy wherein LINGO2-EGFR protein complexes were generated in a cell-free system and co-incubated with either media only, irrelevant cytokine rIL-2 (1ng/ml), or rTFF3 (1ng/ml) followed by immunoprecipitation with LINGO2-Flag and immunoblotted for detection of EGFR-GFP. In support of our model, TFF3 treatment reduced the abundance of LINGO2-EGFR complexes whereas complex formation remained intact following media only or rIL-2 treatment (Figure 2J). Lastly, we noticed the doubling rate of Δ-LINGO2 cells was consistently higher than the MC38 parental line which, if our model was correct, relied upon constitutive activation of EGFR. Indeed, treatment with the EGFR tyrosine kinase inhibitor Gefitinib^TM^ significantly reduced the doubling rate of Δ-LINGO2 cells over a three-day period but had minimal impact on the parental MC38 line (Figure 2K). These experiments all supported the contention that an otherwise constitutive LINGO2-EGFR repressive complex was disrupted by TFF3, allowing EGFR activation through ligand dependent or independent mechanisms and explaining how TFF3 drives EGFR activation without being a direct EGFR ligand ^29^.

Although TFF3KO mice have been reported particularly sensitive to dextran sodium sulfate (DSS)-induced colitis ^6^, our data do not support this contention inasmuch as EGFR inhibition in the TFF3 KO is required to increase their susceptibility to colitic disease, suggesting that LINGO2-dependent inhibition of EGFR signaling helps keep colitis in check. To determine if this effect is mediated through LINGO2, LINGO2KO and co-housed WT littermates were orally administered DSS to induce colitic tissue injury. As reported for DSS-treated TFF3KO mice, LINGO2KO mice had an exacerbated disease phenotype at either 3 or 2.5% DSS marked by significantly greater weight loss, disease activity index (Figure 3A-C), and mortality (Figure S3A) compared to WT controls. Five days following removal of DSS from the drinking water, colon immunopathology was more severe in LINGO2 deficient mice, with greater degree of submucosal edema, ulceration, smooth muscle hyperproliferation and inflammatory infiltration as compared to WT controls (Figure 3D). Consistent with STAT3-mediated protection against colitic disease severity, LINGO2 was necessary for enterocyte expression of pSTAT3, but curiously lamina propria expression of pSTAT3 remained intact (Figure S3B), suggesting a selective role for LINGO2 in the epithelial compartment during the recovery from DSS-induced colitis. Exacerbated colitic disease in LINGO2KO mice was rescued by Gefitinib treatment shown by reduced weight loss (Figure 3E) and increased colon lengths compared to LINGO2KO vehicle treated groups (Figure 3F-G). Thus, the phenotypic consequences of LINGO2 deficiency in the context of DSS colitis were largely due to heightened EGFR activity.

**Figure 3.**
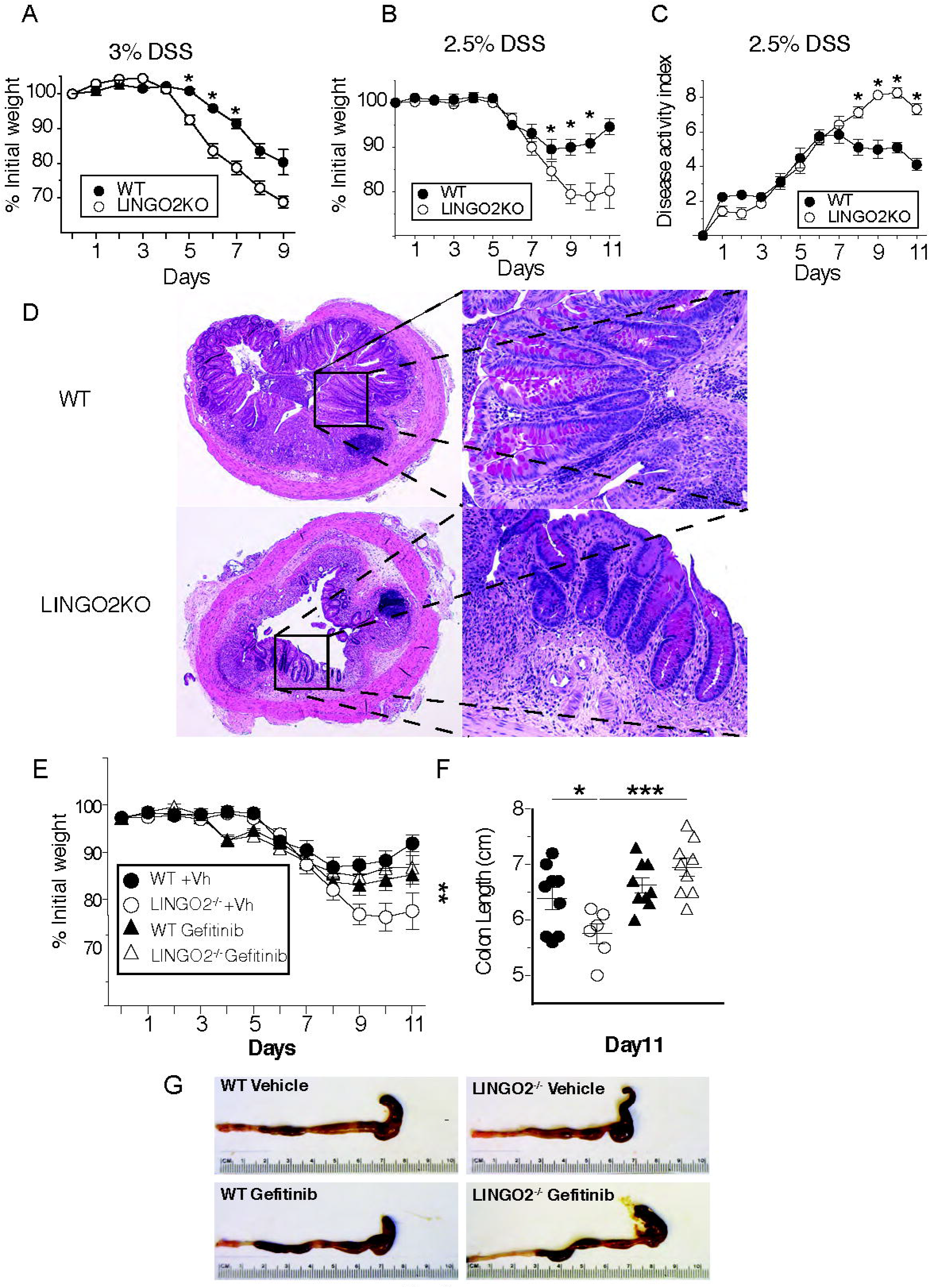
LINGO2/EGFR axis controls DSS colitis severity. (A) Change in initial body weight caused by dextran sodium sulfate administration at 3% or (B) 2.5% w/v in cohorts of WT versus LINGO2KO mice. (C) Disease activity clinical scores in WT versus LINGO2KO mice from mice in “B” Mean ± SE of 8-10 mice/group are shown. (D) H&E and PAS stain for WT (C-D) and LINGO2 KO (E-F) rectum 11 days post 2.5% DSS treatment. Representative images are shown. Magnification 4x (left) and 40 x (right)(E) Change in initial body weight in WT versus LINGO2KO mice post 2.5% DSS treatment with or without Gefitinib. Means ± SE of 8-10 mice/group are shown. (F-G) Colon length in WT versus LINGO2^-/-^ mice at day 11 post 2.5% DSS with or without Gefitinib. Means ± SE of 8-10 mice/group are shown (*\*P < 0.05; **P < 0.01, ***P < 0.001*).

EGFR signaling drives host protection against multiple gastrointestinal nematode species through mechanisms that rely upon T_H_2 cells and ILC2 ^32^. However, intestinal epithelia also have significant roles in host immunity, as intestinal epithelia lineages such as goblet cells secrete many anti-helminth effector molecules including: Muc2, TFF2, Muc5ac, and Relm beta ^33-35^. To determine if TFF3 regulates host immunity against GI nematode infection ^36^, we injected *Nippostrongylus brasiliensis* (*N.b.)* larvae subcutaneously. Following injection, larvae transiently migrate through lung parenchyma, down the esophagus, and into the jejunum within 3 days where they develop into fecund adults that are expelled between 9-12 days post-infection through innate and adaptive Type 2 cytokines (e.g., Areg, interleukins 4, 5, 9, 13, 25 and 33) ^37^ (Figure 4A). Quantitative RT-PCR and immunostaining for TFF3 within the jejunum of naïve and *N.b*.-infected mice demonstrated a marked infection-induced increase of TFF3 positive goblet cells (Figure 4B and Figure S4A). Therefore, we next tested whether TFF3-LINGO2-EGFR de-repression was also operative in the context of parasitic worm infection. *N.b*.-infected TFF3KO mice harbored significantly more adult worms than WT mice at 9 days post infection (Figure 4C). TFF3KO susceptibility did not associate with impaired Type 2 cytokine production but was marked instead by increased pro-inflammatory cytokines such as IFN-γ that impair worm expulsion (Figure 4D). Our model predicted that, if LINGO2 were absent, enhanced EGFR activity would accelerate worm expulsion and, indeed, LINGO2KO mice were highly resistant to *N. b*. infection (Figure 4E) and terminated fecal egg production more rapidly (Figure 4F) than did WT mice. This phenotype was observed regardless of co-housing or the sex of the animals (not shown). Consistent with the presence of fewer parasites, LINGO2KO mice produced less Type 2 cytokine than WT mice (Figure 4G-H), but expressed higher levels of Areg and *Hb-Egf* at day 9 compared to WT littermates, which is consistent with EGFR signaling as a driver of worm clearance ^32^. Irradiation BM chimeras demonstrated that loss of LINGO2 in the nonhematopoietic compartment was responsible for enhanced immunity (Figure 4I). Immunoblotting for pEGFR activity within intestinal lysates revealed that infected LINGO2KO mice had stronger induction of pEGFR compared to WT infected mice (Figure 4J-K) and, consistent with our model’s prediction, inhibition of EGFR signaling following Gefitinib administration reversed the resistant phenotype and resulted in significantly higher adult worm numbers in comparison to vehicle-treated control mice (Figure 4L). Thus, enhanced EGFR activity is responsible, at least in part, for enhanced immunity in LINGO2KO mice.

**Figure 4.**
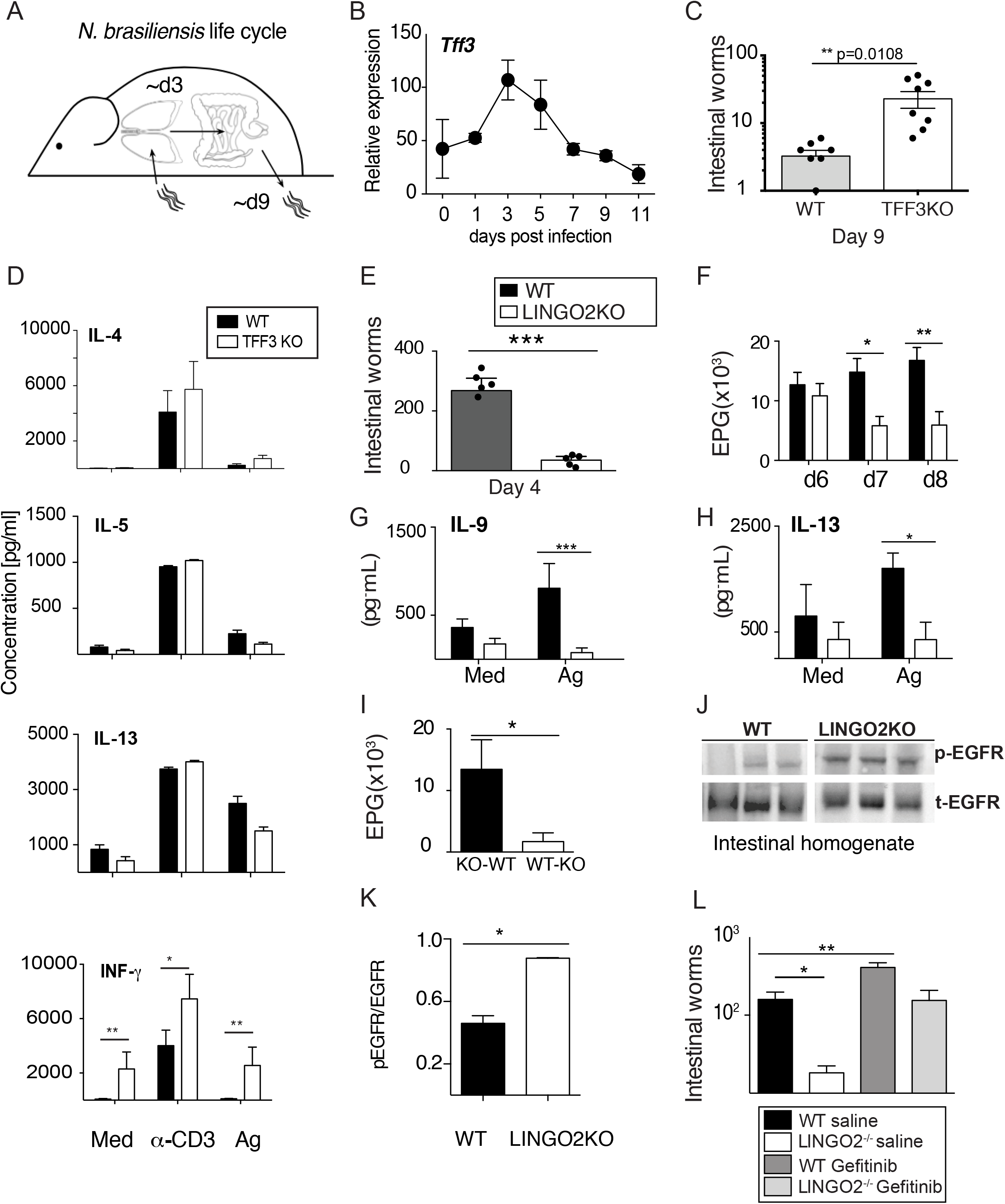
Non-hematopoietic TFF3-LINGO2-EGFR axis controls host-immunity against hookworm infection. (A) Diagram of *N. brasiliensis* life cycle in mice. (B) *Tff3* transcript levels in jejunal mRNA in WT C57BL/6 mice infected with 750 L3 of *N. brasiliensis* Means ± SE of 3-4mice/group. (C) Intestinal adult worm numbers in WT vs. TFF3KO mice at day 9 post-infection each symbol represents an individual mouse. (D) Secretion of IL-4, IL-5, IL-13 and IFN-g levels from WT vs. TFF3KO mesenteric lymph node cells isolated at day 9 post-infection. Cytokine levels in response to media alone, anti-CD3 (1μg/ml or N. d crude adult extract (10μg/ml) at 48hr. (E) Intestinal worm numbers at day 4 post-infection 9 days post infection with *N. brasiliensis* in WT versus LINGO2KO mice. Means ± SE of 5 mice/group are shown (*\*P < 0.05; **P < 0.01, ***P < 0.001*) representative of 4 independent experiments. (F) Fecal egg numbers in WT vs. LINGO2KO mice at days indicated. (G) IL-9 and (H) and IL-13 production from isolated MLN on day 9 exposed to media or *N. brasiliensis* antigen extract. **(I)** Egg load in feces on day 7 from WT→LINGO2^-/-^ versus LINGO2^-/-^→WT BM chimera mice at day 4 post infection Means ± SE of 3-4 mice/group are shown (*\*P < 0.05*). (J) Western blot showing intestinal levels of phospho-EGFR and total EGFR in mice at day 9 post-infection Each represents an individual mouse (K) Densitometry for data shown in “J” (L) Adult worm burdens recovered from intestinal lumen at day 9 from WT and LINGO2^-/-^ mice treated with Gefitinib or vehicle during infection with *N. brasiliensis*. Means ± SE of 9-10 mice/group are shown (*\*P < 0.05; **P < 0.01*).

In conclusion, we propose a previously-unappreciated mechanism wherein TFF3, a reparative cytokine of the mucosal interface, binds to LINGO2 and thereby releases tonic inhibition of EGFR activity. This de-repression mediated by TFF3 then regulates the extent of EGFR activity similar to a rheostat: in the absence of LINGO2, excessive EGFR activity in epithelia leads to increased baseline proliferation and greater colitic disease severity whereas, in the context of worm infection, it enhances IEC-driven host immunity. In both cases, reduction of EGFR activity reverses these phenotypes. It is likely that this TFF3-LINGO2-EGFR axis has distinct roles in hematopoietic vs. non-hematopoietic cell lineages and may be pertinent to other disease states involving autoimmunity and tumorigenesis.

## Methods

### Mice

All animal procedures were approved by the Institutional Animal Care and Use Committee at University of California, San Francisco. Super-ovulated female C57BL/6 mice (4 weeks old) were mated to C57BL/6 stud males. Fertilized zygotes were collected from oviducts and injected with Cas9 protein (30 ng/ul), sgRNA (15 ng/ul) into pronucleus of fertilized zygotes. Injected zygotes were implanted into oviducts of pseudopregnant CD1 female mice. Animals were housed under specific-pathogen free barriers in vivarium at San Francisco General Hospital or University of Pennsylvania. All the procedures were reviewed and approved by IACUC at University of California at San Francisco (protocol #AN109782-01) and University of Pennsylvania (protocol #805911).

### DSS colitis and Nippostrongylus brasiliensis mouse models

Colitis was induced in mice using 2.5% DSS (w/v, 0216011080, MP-Biomedicals LLC Solon, OH) in autoclaved drinking water for 6 days. Normal water was supplied on Day 6. Mice were assessed for body weight, appearance, fecal occult blood (Fisherbrand™ Sure-Vue™ Fecal Occult Blood Slide Test System, Thermo Fisher Scientific, Inc., Waltham, MA, USA), stool consistency, and diarrhea during the entire course of colitis. Colon length was determined using a ruler at necropsy. Disease activity index (DAI) a cumulative score with a maximum of 12, was derived as follows: every mouse was assigned a score of 1 Vs 0 was for each display of lack of grooming, piloerection, awkward gait, hunched posture, and lack of mobility. Diarrhea score (0, Normal stool, 1 soft stool, 2 loose formed stools and 3, watery fecal matter), Rectal Bleeding was scored as 0 no blood, 1 minor bleeding and 2 gross rectal bleeding). Fecal occult blood test was scored as 0, Negative, 1 Mild amount of blood and 2 excessive presence of blood. Pharmacological inhibition of EGFR was achieved by administering Gefitinib (10mg/Kg body weight i.p., G-4408 LC Laboratories, Woburn, MA) in DMSO. Parasites were maintained on fecal culture incubated at 25C. Larvae from 7 to 14-day old cultures were collected and washed 3x in PBS with 1% Penicillin/Streptomycin. To infect, 650-700 L3 stage larvae per mouse were injected subcutaneously to the base of tail. On d6-8 fecal pellets were collected from each mouse to assess egg counts. On d9, mice were euthanized and the proximal half of small intestines were removed, opened longitudinally, and incubated in PBS for 2h before collecting and counting adult worms.

### Cloning/plasmid construction

The full-length mLingo2 coding region was obtained with KpnI and XhoI double digestion of Lingo2 cDNA plasmid (MR215769, Origene) and subcloned into pcDNA3:GFP vector (Addgene) or pcDNA3:mRFP vector (Addgene) to generate pcDNA3:Lingo2:GFP and pcDNA3:Lingo2:RFP constructs. A serial of deletion of Lingo2 functional domains (Ecto and cytoplasm tails) were generated by PCR amplification with the specific primers bearing KpnI and XhoI overhangs, and cloned into pcDNA3:GFP vector (Adgene) to get Lingo2 truncated constructs fused with GFP. The full-length mTFF3 coding region was amplified with mTFF3 forward primer with HindIII overhang and mTFF3 reverse primer with XhoI overhang using mTFF3 cDNA plasmid (MR200151, Origene) as the template. The digested TFF3 PCR product with HindIII and Xho enzymes was cloned into pcDNA3:mRFP vector (Addgene) to generate pcDNA3:mTFF3:RFP construct. Using the similar method, the full-length mTFF3 coding region was also cloned into pINFUSE-mIgG2b-Fc2 vector (InvivoGen) to generate pINFUSE-TFF3:mIgG2b-Fc2 construct. The full-length mCXCR4 coding region was amplified with mCXCR4 forward primer with HindIII overhang and mCXCR4 reverse primer with XhoI overhang using mCXCR4 cDNA plasmid (OMu23034D, Origene) as the template. The digested CXCR4 PCR product with HindIII and Xho enzymes was cloned into pcDNA3:GFP vector (Addgene) to generate pcDNA3:mCXCR4:GFP construct. The full-length SDF1/CXCR12 coding region was amplified with mSDF1 forward primer with EcoRI overhang and mSDF-1 reverse primer with XhoI overhang using SDF1 cDNA plasmid (MR227229, Origene) as the template. The digested SDF1 PCR product with EcoRI and Xho enzymes was cloned into the pcDNA3:mRFP vector (Addgene) to generate pcDNA3:SDF1:RFP construct. The sequences of constructs were confirmed by DNA sequencing in UPenn Sequencing facility.

### Human tissue and immunostaining

Archived rectosigmoid biopsy samples were from the SCOPE cohort at the University of California, San Francisco (UCSF). The SCOPE cohort is an ongoing longitudinal study of over 1,500 HIV-infected and uninfected adults followed for research purposes. The UCSF Committee on Human Research reviewed and approved the SCOPE study (IRB# 10-01218), and all participants provided written informed consent. PLA starter Duolink^®^ In Situ Orange Starter Kit Mouse/Goat was obtained from Sigma Aldrich containing the PLA probes anti-goat MINUS and anti-mouse PLUS with the detection reagent DuoLink^®^ Orange. Fluorescence for proximity ligation was observed under a Cy3 filter on a Leica Inverted Microscope DMi8 S Platform. The primary antibodies for anti-LINGO2 Ab was raised against Goat (R&D Systems; AF3679) and anti-TFF3 raised against mouse (eBioscience; 5uclnt3) were used to detect the LINGO2 and TFF3 interactions in situ. Technical controls omitting either one of the antibodies served as a negative control as no circular DNA oligonucleotides were able to be amplified. Formalin fixed paraffin embedded tissue was stained for primary antibodies based on established staining protocols.

### Transfection and immunofluorescence

HEK293 cells were maintained in DMEM medium (Invitrogen) supplemented with 10% fetal bovine serum (Invitrogen) and 1% penicillin-streptomycin (Invitrogen) at 25°C with 5% CO2. Before transfection, 2 ml of cells (0.2 × 106/ml) per well was seeded on in a well of Lab-Tek Chamber slide (sigma) for overnight culture. 1ug plasmids Lingo2:GFP and TFF3:RFP were co-transfected to the cells using X-tremeGENE HP DNA transfection reagent (Roche) based on the manufacture’s manual. For colocalization of EGFR:GFP and Lingo2:RFP, pCMV3-mEGFR-GFPSpark(MG51091-ACG, Sino Biological Inc.) and pDNA3-Lingo2:RFP was transfected in the same method. After three-days culture, the cells were fixed with 4% paraformaldehyde in PBS for 10 min and stained with DAPI (0.5ug/ml) for 5 mins. After mounting the cells with the coverslip, Lingo2:GFP and TFF3:RFP were visualized using the Leica TCS-NT confocal microscope.

### Co-Immunoprecipitation

HEK293 or CHO cells were transfected with 2 ug pCMV6:Lingo2:Flag (MR215769, Origene) plasmid. After incubation of the transfection mixture with the cells for 72 h, the cells were washed twice with PBS and lysed with 0.5 ml Digitonin extraction buffer [1% digitonin; 150 mM NaCl; 1 mM MgCl2;10 mM TrisCl, pH 7.4, 1X Proteinase Inhibitor (Sigma)]. The protein concentration was be measured using BCA assay kit (Invitrogen). Around 0.6 mg total protein was incubated with 10μg TFF3-Fc (Ray Biotech) or Fc (Ray Biotech) overnight and precipitated with 30μl protein G agarose (Invitrogen) for 2□h at 4°C. The IP complex were washed with PBS for three times and eluted with 2× SDS loading buffer at 95°C. The IP complex along with 5% input were immunoblotted with mouse anti-Flag antibody (1:5000) (Sigma) to detect the interaction between Lingo2:Flag and TFF3-Fc. LINGO2 was detected by rabbit anti-FLAG followed by incubation with goat anti-rabbit IgG Cy5 and goat anti-mouse IgG Cy3 antibodies. To detect EGFR and Lingo2 interactions, HEK293 were cotransfected with 2 ug pEGFR:Flag (MG51091-CF, Sino Biological Inc.) and 2 ug pDNA3:Lingo2:GFP plasmids as described above. The cells were cultured for 72 h and extracted with digitonin for the cell lysis. Around 0.5 mg total protein was precipitated using 2μg normal rabbit IgG (the negative control), 2ug anti-GFP rabbit polyclonal antibody (Torrey Pines Biolabs), 2μg normal mouse IgG (the negative control), 2ug anti-flag mouse antibody (Sigma) overnight and further incubated with 30μl protein A agarose (Invitrogen) for 2□h at 4°C. The IP complex precipitated with anti-GFP antibody along with 5% input was immunoblotted with mouse anti-Flag antibody (1:5000) (Sigma) to detect the interaction between Lingo2:GFP and EGFR-flag. The IP complex precipitated with anti-flag antibody along with 5% input was immunoblotted with rabbit anti-GFP antibody (1:5000) for the detection of reciprocal interaction. For determining if TFF3 disrupts the interaction between Lingo2 and EGFR, the mixture of 100ug Lingo2:GFP lysate and 50ug EGFR-flag lysate was incubated with either 2μg TFF3 or 2μg mIL2 (R& D System) proteins for two hours at 4°C. Then, Co-IP was performed with either rabbit anti-GFP antibody or normal rabbit IgG as above. The IP complex was immunoblotted with mouse anti-Flag antibody (1:5000) (Sigma) to detect the interaction between Lingo2:GFP and EGFR-flag. IL-10 measurement by ELISA was completed according to the manufacturer’s protocol.

### Generation of a Lingo2 knockout cell line

LINGO2 Knockout colon epithelial cell line (MC38) was generated based on HR (homologous recombination)-mediated CRISPR–Cas9 genome editing. Validated LINGO2 guide RNA obtained commercially (PNA Bio). Briefly, MC38 cells were maintained in DMEM medium (Invitrogen) supplemented with 10% fetal bovine serum (Invitrogen), 1XNEAA, 10mM HEPES, 1mM Sodium Pyruvate, 50ug/ml Gentamicin and 1% penicillin-streptomycin (Invitrogen) at 25°C with 5% CO2. Before transfection, 3 ml of cells (1 × 105/ml) per well was seeded in a well of a six-well plate for overnight culture. 1ug Lingo2 sgRNA, 6ug Lingo2 donor vector and 1ug Cas9 plasmid were co-transfected into the cells using X-tremeGENE HP DNA transfection reagent (Roche). After expanding the cells for the two generations, H2KK positive cell (LINGO2KO cells were identified by flow cytometry because Lingo2 donor vector contains a H2KK gene cassette. Loss of Lingo2 expression in H2KK positive cells was further confirmed by qRT-PCR analysis.

Western blotting: 1 ml of M38 parental or Ling2 KO cell (1 × 105/ml) per well was seeded in a well of a 24-well plate and cultured for 48 hrs in complete medium. The cells were washed with 1ml PBS and cultured in 1ml serum-free medium for 24 hrs for serum starvation. Then, the cells were treated with TFF3-Fc (1ug/ml) in the complete medium for 0, 5min, 15min, 30min and 1hr. After washing the cells with 1ml PBS twice at the indicated time, the cells were lysed with the lysis buffer [1% Triton X-100, 150 mM NaCl,10 mM Tris (pH 7.4), 1 mM EDTA, protease and phosphatase inhibitor cocktail (Sigma)]. The mouse colon tissues from WT or Lingo2 KO mice were lysed with RIPA buffer containing protease and phosphatase inhibitor cocktail (Sigma). 15ug samples were loaded and separated in 4-12% PAGE gel (Invitrogen). The following primary Abs were used for immunoblotting assays: mouse anti-Stat3 (Cell signaling,1:5000), rabbit anti-pStat3(Y705) (cell signaling, 1:1000), rabbit anti-EGFR (Millerpore,1:1000), rabbit anti-pEGFR (Y1068) (Abcam,1:1000); mouse anti-actin (Santa Cruz, 1:2500).

### TRICEPS Mass spectrometry

The CaptiRec samples were analyzed on a Thermo LTQ Orbitrap XL spectrometer fitted with an electrospray ion source. The samples were measured in data dependent acquisition mode in a 40 min gradient using a 10cm C18. The 12 individual samples in the CaptiRec dataset were analyzed with a statistical ANOVA model. This model assumes that the measurement error follows Gaussian distribution and views individual features as replicates of a protein’s abundance and explicitly accounts for this redundancy. It tests each protein for differential abundance in all pairwise comparisons of ligand and control samples and reports the p-values. Next, p-values are adjusted for multiple comparisons to control the experiment-wide false discovery rate (FDR). The adjusted p-value obtained for every protein is plotted against the magnitude of the fold enrichment between the two experimental conditions. The area in the volcano plot that is limited by an enrichment factor of 2-fold or greater and an FDR-adjusted p-value less than or equal to 0.01 is defined as the receptor candidate space. To allow for statistical analysis, the experiment was done in biochemical triplicates. The samples enriched in glycopeptides were analyzed on a Thermo LTQ Orbitrap XL spectrometer. Peptide identifications were filtered to a false-discovery rate of <= 1% and quantified using an MS1-based label-free approach.

### Histology and colon biopsies

Mice were eviscerated, and colons extracted. A small biopsy of the intact distal colon was excised, fixed overnight at 4°C in neutral 10% formaldehyde, further dehydrated and embedded in paraffin for histological analysis. Another small biopsy in the distal region was snap frozen in liquid N2 for protein analysis. The remaining colon was opened longitudinally, flushed with ice-cold PBS to remove feces, six punch biopsies were obtained at the proximal end, the remaining perforated regions were stored for 1h in RNAlater (AM7021, Thermo Fisher) on ice and subsequently stored at −80°C until use. Remaining length of the colon was embedded as “swiss rolls” in OCT. Pathological assessment of colitis was performed blindly by certified pathologists as per previously described criteria.1-3

### Immuno-histochemistry and Imaging

Staining for pSTAT3 was performed as previously described,4 briefly, 5μm thick paraffin embedded sections were dewaxed though descending grades of Ethanol to water. Heat induced antigen recovery was further performed on the hydrated sections by boiling the slides in 1mM EDTA in a pressure cooker for 1h. Sections were further permeabilized with TBS containing 0.2% Triton X-100 (X100, Sigma-Aldrich, St. Louis, MO), all sections were then incubated with 2%BSA containing 10% Goat Serum in TBS to block non-specificity under humid conditions. Sections were then incubated overnight with antibody against pTyr705-STAT3 (9145, Cell Signaling, Danvers, MA) and isotype control (500-P00, Peprotech, Rocky Hill, NJ). Next day the sections were rinsed times in TBS containing 0.1% Tween 20 (BP337-500, ThermoFisher) and further incubated with HRP conjugated secondary Ab, at RT (SignalStain^®^ Boost, Cell Signaling 8114), for 40mins under humid conditions. Sections were rinsed with TBS-0.1% Tween 20 and further with TBS. Bound antibody was detected by treating the sections with ImmPACT DAB Peroxidase Substrate (SK-4105, Burlingame, CA) as per manufacturer’s instructions and further counterstained with hematoxylin (26306-01, Electron Microscopy Sciences, Hatfield, PA). Images were acquired using a Leica DM6000B Upright Widefield Microscope and further analyzed using MetaMorph image analysis software (Molecular Devices, LLC, San Jose, CA). Nuclear pSTAT3 intensity was calculated using MetaMorph software in defined areas of interest on epithelial cells and lamina-propria cells per high power field. Intensities were assigned by MetaMorph on a color scale of (0 – 255) with 0 denoting Black, and 255 White respectively, i.e. lower value denoting a more intense/darker staining and vice-versa.

#### RNA isolation and real-time qPCR

Approximately 0.5 cm of duodenum was removed for RNA extraction using NucleoSpin RNA Plus kit (Macherey-Negel, Dueren, Germany). Five hundred nano-grams of total RNA were used to generate cDNA with Super Script II (Invitrogen). Quantitative real-time PCR was performed on the CFX96 platform (Bio-Rad, Hercules, CA). Gene expression levels were normalized to (*Gapdh*).

#### Statistics

Statistical analyses were performed using GraphPad Prism version 7.0 (GraphPad, La Jolla, CA). A two-tailed Student’s T test or ANOVA was used where appropriate.

## Author Contributions

Y.J., Y.W., J.P., LY.H., T.Y., K.H., T.O., C.P., W.N., conducted experiments, M.S., contributed valuable reagents and samples, Y.J., Y.W., J.P., helped write the manuscript and D.R.H., conceived the study and wrote the manuscript.

